# Single-nucleus RNA-seq atlas of 5xFAD mouse brain maps cell-type-specific transcriptional disruption and preferential rolipram rescue in VIP interneurons

**DOI:** 10.64898/2026.11.11.765944

**Authors:** Augustina Potokiri, Abena Asua Dwamena, Hongmin Wang

## Abstract

**Background:** Alzheimer’s disease is characterized by progressive synaptic dysfunction and inhibitory circuit disruption across multiple brain cell types. Rolipram, a selective phosphodiesterase 4 (PDE4) inhibitor that elevates intracellular cyclic AMP (cAMP), activates proteasome, and produces robust procognitive effects in AD mouse models. Still, the cell-type-specific transcriptional mechanisms underlying these effects remain uncharacterized.

**Methods:** We profiled 29,664 nuclei from 5xFAD and wild-type (WT) mouse right hemisphere across four conditions: WT Vehicle, WT Rolipram, 5xFAD Vehicle, and 5xFAD Rolipram (0.03 mg/kg/day, 30-day subcutaneous osmotic pump) using the Parse Biosciences Evercode WT v3 combinatorial barcoding platform. Nuclei were clustered into 16 annotated cell type populations using Seurat v5. Disease and rescue effects were quantified as log fold changes (LFCs) across all clusters, with Gene Ontology enrichment assessed using clusterProfiler and transcription factor activity inferred using decoupleR.

**Results:** Disease-associated transcriptional disruption was highly cell-type-specific in its magnitude. Among inhibitory subtypes, VIP/Reln disinhibitory interneurons (n = 149 nuclei) showed the most coherent transcriptional disruption: 289 candidate genes were lower in 5xFAD, with Gene Ontology enrichment concentrated in synapse organization programs, including regulation of synapse structure or activity (padj = 1.5×10^−14, 44 genes), regulation of synapse organization (padj = 1.9×10^−14, 43 genes), and postsynapse organization (padj = 1.1×10^−13, 38 genes). This pathway-level specificity was not reproduced in PVALB or SST interneurons. Astrocytes showed loss of cell-adhesion programs (cell-substrate adhesion padj = 2.4×10^−4, 10 genes). Rolipram rescue was preferential: VIP interneurons yielded 72 true rescue genes and astrocytes yielded 42, with the remaining 14 clusters showing substantially fewer. The cAMP-specific phosphodiesterase Pde7b was disease-context-specifically lower in 5xFAD VIP interneurons (LFC −0.53) and restored by rolipram (LFC +0.54), with WT expression unchanged. decoupleR CREB1 activity inference identified reduced CREB1 activity in 5xFAD VIP interneurons with partial restoration in 5xFAD Rolipram, supporting a stepwise transcriptional cascade in which amyloid pathology reduces cAMP, which suppresses PKA and CREB1 activity. This in turn reduces Pde7b transcription, and rolipram reverses this cascade by blocking cAMP hydrolysis upstream.

**Conclusion:** This single-nucleus atlas provides the first cell-type-resolved characterization of rolipram’s transcriptional effects in the 5xFAD brain. VIP disinhibitory interneurons emerge as the most transcriptionally vulnerable inhibitory subtype and the preferential site of rolipram-associated rescue, with the Pde7b/CREB1 signaling axis nominating a tractable, disease-context-specific pharmacological substrate.

## Introduction

Alzheimer’s disease (AD) is the leading cause of dementia worldwide, affecting approximately 6.9 million Americans aged 65 and older and an estimated 55 million individuals globally, with prevalence projected to nearly triple by 2060 as the population ages (1). The disease is characterized by progressive cognitive decline driven by amyloid-beta (Abeta) plaque deposition, neurofibrillary tau tangles, synaptic loss, and neuroinflammation. The recent approvals of lecanemab and donanemab, monoclonal antibodies targeting aggregated Abeta, have validated amyloid as a clinically relevant therapeutic target. In the phase 3 CLARITY-AD trial, lecanemab significantly reduced amyloid burden and slowed clinical decline by approximately 27% over 18 months compared with placebo (2). Yet amyloid clearance alone does not stop disease progression or restore established synaptic dysfunction. The downstream, cell-type-specific molecular mechanisms that drive cognitive loss remain incompletely characterized and identifying them matters precisely because synaptic circuitry can still be rescued at earlier disease stages, making those mechanisms more accessible targets for intervention than amyloid itself.

The modest clinical benefits achieved by current amyloid-targeting therapies highlight an important conceptual gap: plaque clearance does not reverse the synaptic and circuit-level dysfunction that underlies cognitive impairment once it is established. Soluble amyloid-beta oligomers disrupt synaptic transmission, impair long-term potentiation, and suppress the network activity patterns required for memory encoding at a stage that precedes detectable neurodegeneration (11). These oligomer-driven effects are mediated at the level of individual synapses but propagate through the inhibitory microcircuits that regulate cortical gain, attention gating, and memory consolidation. Effective disease modification therefore requires not only amyloid clearance but concurrent restoration of the synaptic and circuit-level programs that amyloid pathology disrupts. Defining which cell populations and molecular pathways are functionally compromised and pharmacologically rescuable is a foundational step toward rational, mechanism-based therapeutic development in AD.

High-throughput single-nucleus RNA sequencing (snRNA-seq) has fundamentally transformed our ability to dissect cellular heterogeneity in the AD brain. Landmark studies in human post-mortem tissue have mapped cell-type-specific transcriptional changes at single-cell resolution, revealing a homeostatic-to-disease-associated microglia (DAM) transition in microglia (3, 4, 5), reactive astrogliosis programs associated with neuronal injury (6), and interneuron-specific synaptic gene losses in excitatory and inhibitory neurons (7). The scale of such profiling has expanded dramatically: a cell atlas of approximately 2.3 million single nuclei from 427 individuals in the ROSMAP cohort identified transcriptional correlates of cognitive resilience and dementia, emphasizing that cell-type-specific programs in neurons and glia together determine disease trajectory (8). A subsequent multiregion atlas spanning six brain regions confirmed that transcriptional disruption is not restricted to neurons but encompasses coordinated pathological programs in microglia, astrocytes, oligodendrocytes, and oligodendrocyte precursor cells (OPCs) (7). Complementary multimodal profiling in human cortex combining chromatin accessibility with transcriptomics has further shown that cell-type-specific regulatory networks are reorganized in AD (9), and paired human-mouse snRNA-seq has established the degree to which rodent model transcriptomes recapitulate key human disease signatures, including TREM2-dependent microglial responses (10). When a transcriptional program is lost selectively within a defined interneuron subtype, the circuit functions of that subtype become primary candidates for the underlying pathophysiological mechanism.

The 5xFAD transgenic mouse model carries five familial AD mutations: the Swedish (K670N/M671L), Florida (I716V), and London (V717I) mutations in human amyloid precursor protein (APP), and the M146L and L286V mutations in presenilin-1 (PSEN1). Together, these mutations drive robust intraneuronal Abeta accumulation and extracellular plaque deposition detectable as early as 2 months of age (12, 13). By 6-9 months of age, the window used in the present study, 5xFAD mice exhibit extensive cortical and hippocampal amyloid burden, progressive synaptic protein loss, selective neurodegeneration in cortical layer 5, and deficits in spatial memory and contextual fear conditioning, establishing the model as one of the most widely used platforms for preclinical therapeutic evaluation. Despite this extensive behavioral and neuropathological characterization, the cell-type-specific transcriptional landscape of the 5xFAD cortex, particularly within inhibitory interneuron subtypes, and its modulation by pharmacological intervention have not been resolved at single-nucleus resolution.

A mechanistically undercharacterized dimension of AD circuit pathophysiology is the selective disruption of cortical inhibitory interneuron networks. The cortical GABAergic system comprises molecularly and functionally distinct subtypes organized into hierarchical inhibitory microcircuits. VIP-expressing interneurons occupy a specialized disinhibitory position within cortical microcircuits, providing preferential GABAergic input onto SST-positive interneurons that exert dendritic inhibition over pyramidal neurons; VIP interneuron activation therefore produces net disinhibition of excitatory pyramidal output through an inhibition-of-inhibition motif (14, 15, 16). This disinhibitory microcircuit gates cortical gain during behavioral states of heightened attentional arousal and cholinergic/noradrenergic neuromodulatory drive, precisely the conditions that permissively regulate Hebbian synaptic plasticity and the induction of long-term potentiation. Proteomic profiling in 5xFAD mice has revealed that parvalbumin (PV) interneurons exhibit early presynaptic and postsynaptic vulnerability, with progressive loss of active zone scaffolding and postsynaptic density proteins detectable at 3 months of age, prior to the emergence of bulk transcriptomic changes (17). Transcriptomic profiling of PV- and SST-positive interneurons in 5xFAD hippocampus and cortex has further revealed region- and subtype-restricted gene expression dysregulation that evolves with disease stage, implicating impaired oxidative phosphorylation, disrupted GABAergic synaptic transmission, and interneuron-autonomous neuroinflammatory signaling as convergent pathological mechanisms (18). VIP interneurons show altered intrinsic excitability and CA1 disinhibitory circuit dysfunction in the 3xTg-AD model, with VIP/calretinin-coexpressing cells displaying measurable changes in action potential threshold, amplitude, and afterhyperpolarization kinetics at a stage preceding overt amyloid deposition (19). Together, these findings identify interneuron subtype-selective vulnerability as an early and cell-autonomous feature of amyloid pathology.

Rolipram is a selective inhibitor of phosphodiesterase 4 (PDE4), the predominant cAMP-hydrolyzing enzyme in neurons and glia. PDE4 inhibition elevates intracellular cAMP, thereby activating protein kinase A (PKA). The cAMP-PKA signaling not only activates proteasome activity to enhance tau removal (20) but also drives phosphorylation of the cAMP response element-binding protein (CREB) at Ser133. CREB-mediated transcription is an obligatory effector of late-phase long-term potentiation (L-LTP), activity-dependent synaptic consolidation, and long-term memory formation, and CREB phosphorylation is deficient in both post-mortem human AD cortex and in multiple AD transgenic mouse models (21, 22). Landmark studies established that persistent rolipram treatment rescued synaptic transmission, LTP, and multiple memory tasks in APP/PS1 mice without altering amyloid burden, with cognitive benefits persisting weeks after drug clearance, attributable to restored CREB-dependent gene expression (21). Rolipram also reverses Abeta-peptide-induced memory deficits in rats and reduces neuroinflammation and apoptotic signaling in multiple amyloid-injury models (23). Despite these well-characterized procognitive and synaptotrophic effects, the cell-type-specific transcriptional programs through which PDE4 inhibition restores cognitive function have not been resolved at single-nucleus resolution, leaving unanswered the question of which cell populations serve as the primary transcriptional effectors of cAMP-PKA-CREB signaling restoration in the AD brain.

In this study, we performed snRNA-seq on 5xFAD and compared with wild-type (WT) mice. We provide the first cell-type-resolved characterization of rolipram’s transcriptional effects in the 5xFAD brain and demonstrate that VIP/Reln disinhibitory interneurons are the preferential transcriptional responders to PDE4 inhibition, yielding 72 true rescue genes including the cAMP-specific phosphodiesterase Pde7b. Transcription factor activity inference using decoupleR (24) with the CollecTRI CREB1 target gene network identifies reduced CREB1 transcriptional activity in 5xFAD VIP interneurons that is partially restored by rolipram treatment, supporting a cAMP-PKA-CREB1-Pde7b regulatory axis as a disease-context-specific mechanistic candidate.

## Results

### A single-nucleus transcriptomic atlas of 5xFAD and wild-type mouse brain

To characterize cell-type-specific transcriptional changes in 5xFAD brain and their modulation by rolipram we performed snRNA-seq on whole right-hemisphere tissue from 5xFAD and wild-type (WT) mice receiving either rolipram (0.03 mg/kg/day, individually weight-adjusted) or vehicle for 30 days via subcutaneous Alzet osmotic pump implantation (Figure 1A). Following quality control processing through the Parse Biosciences Trailmaker platform 29,664 high-quality nuclei were retained across four experimental conditions: WT Vehicle (n = 8,811), WT Rolipram (n = 7,100), 5xFAD Vehicle (n = 8,840) and 5xFAD Rolipram (n = 4,913). Each condition was represented by a single pooled biological sample comprising whole right-hemisphere tissue from one male and one female animal aged 6.5-8 months at pump implantation and harvested at 7.5-9 months.

**Figure 1.**
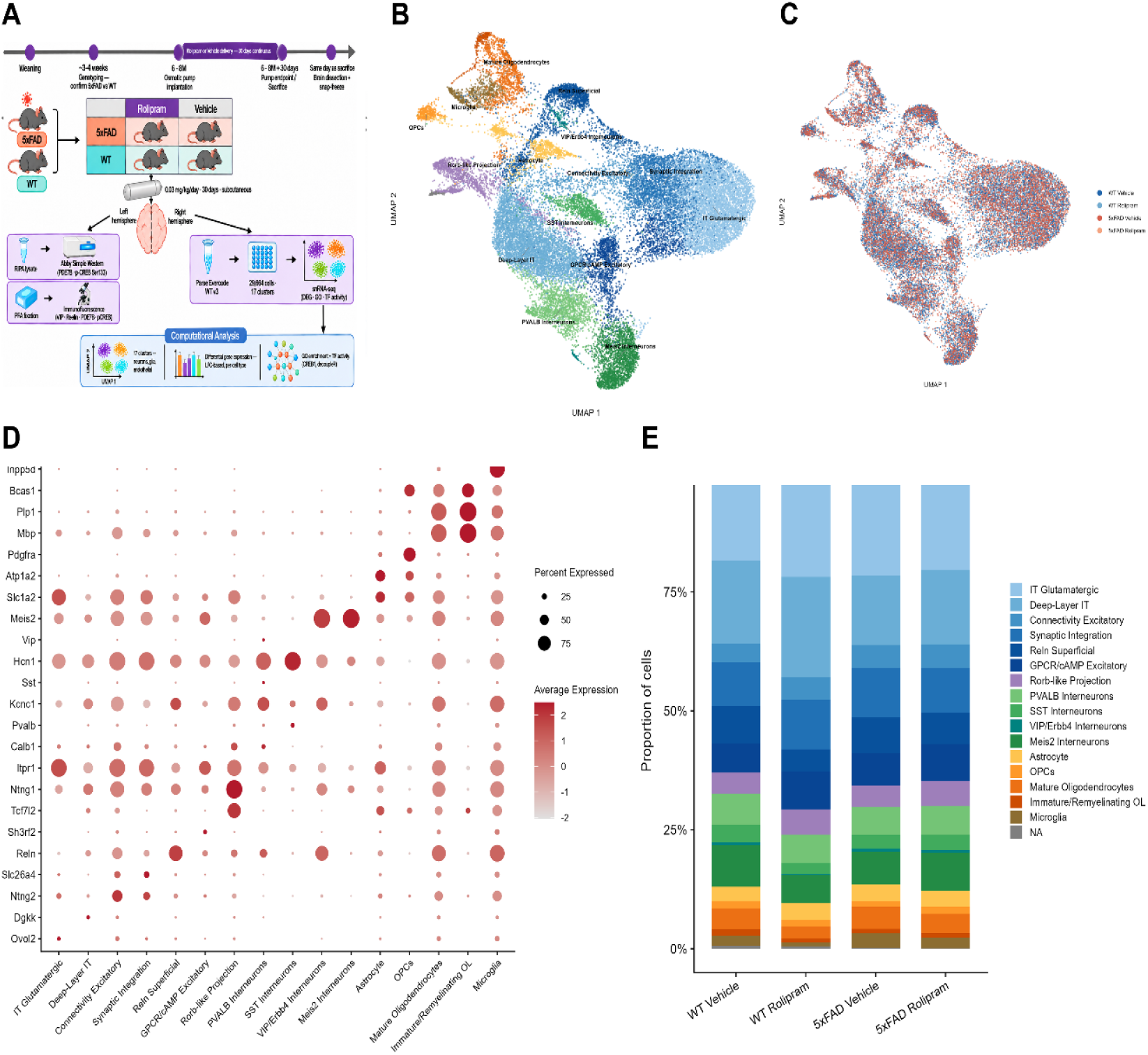
Single-nucleus transcriptomic atlas of 5xFAD and wild-type mouse brain. (A) BioRender schematic of the 2×2 factorial experimental design illustrating the surgical timeline for subcutaneous Alzet osmotic pump implantation, brain harvest and hemisphere assignment, OptiPrep discontinuous density gradient nuclei isolation, Parse Biosciences Evercode WT v3 split-pool combinatorial barcoding library preparation, sequencing and downstream bioinformatic analysis. (B) UMAP projection of 29,664 nuclei colored by annotated cluster identity spanning seven cortical excitatory subtypes, one thalamic excitatory population (Tcf7l2+/Ntng1+), four GABAergic interneuron subtypes (PVALB, SST, VIP/Reln, Meis2) and non-neuronal populations. (C) Same UMAP embedding colored by experimental condition confirming representation of all four conditions across all 16 cluster regions. (D) Dot plot of canonical marker gene expression per cluster where dot size encodes the percentage of nuclei with detectable expression (minimum 1%) and dot color encodes scaled mean expression. (E) Stacked bar chart of proportional cell type composition per experimental group showing broadly stable population fractions across all four conditions.

Unsupervised graph-based clustering resolved 24 transcriptional clusters subsequently consolidated to 16 biologically annotated cell type populations by merging clusters sharing identical marker gene profiles (Figure 1B). The 16 populations comprised seven cortical excitatory neuron subtypes, one thalamic excitatory neuron population, four GABAergic interneuron subtypes (PVALB, SST, VIP/Reln and Meis2), one astrocyte population, two oligodendrocyte lineage populations, one OPC population and one microglial population. Cell type identities were validated by canonical marker gene expression (Figure 1D) cross-referenced against published mouse cortex atlases (26, 27). All 16 clusters were represented across the UMAP embedding (Figure 1C) and cell type proportions were broadly stable across WT Vehicle, WT Rolipram and 5xFAD Vehicle conditions (Figure 1E). One excitatory population was classified as thalamic rather than cortical in origin based on selective enrichment of Tcf7l2 and Ntng1 alongside preferential expression of the vesicular glutamate transporter Slc17a6 (Vglut2) over Slc17a7 (Vglut1) relative to all cortical excitatory clusters; this population is designated Thalamic Excitatory (Tcf7l2+) throughout. The absence of hippocampal and striatal transcriptional signatures indicates the atlas is predominantly cortical with a minor thalamic contribution. The 5xFAD Rolipram condition yielded a lower nuclear yield (n = 4,913) relative to the remaining three conditions (n = 7,100-8,840) which reduces statistical power for rescue analyses in low-abundance clusters and should be taken into account when interpreting cross-cluster rescue comparisons.

### 5xFAD brain shows highly cell-type-specific transcriptional disruption

Computing log fold changes (LFC) between 5xFAD Vehicle and WT Vehicle within each cluster revealed markedly cell-type-specific transcriptional disruption (Figure 2A). The number of candidates differentially expressed genes (|LFC| > 0.25, expressed in >5% of nuclei) ranged from over 400 in astrocytes to fewer than 50 in several excitatory subtypes, establishing that the transcriptional burden of amyloid pathology is nonuniformly distributed across the cortical cellular landscape.

**Figure 2.**
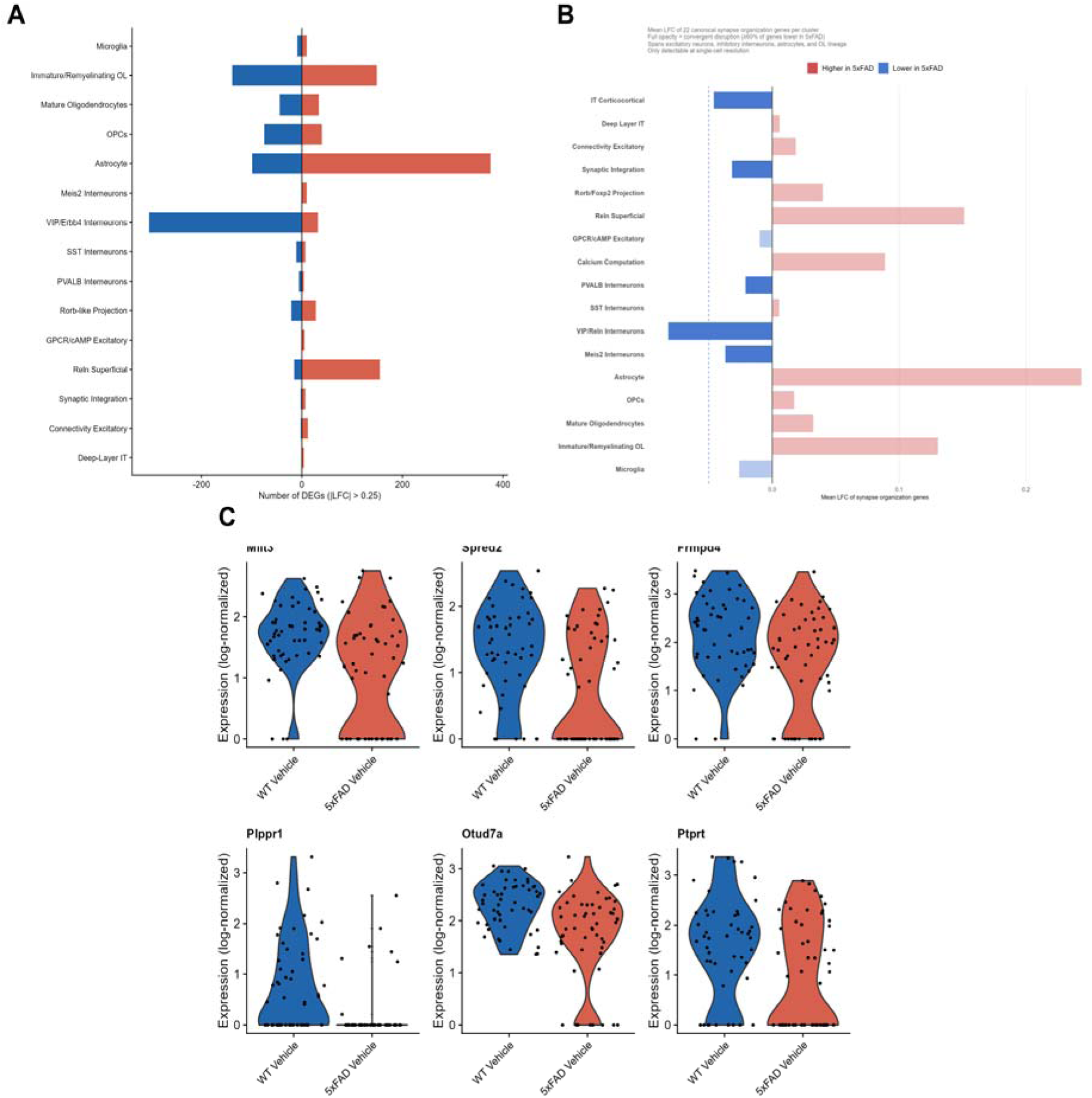
Cell-type-specific transcriptional disruption in 5xFAD brain. (A) Diverging bar chart of candidate differentially expressed gene counts per cluster (5xFAD Vehicle vs WT Vehicle, |LFC| > 0.25, >5% nuclei) demonstrating nonuniform transcriptional burden across the cortical cellular landscape. (B) Diverging bar chart of mean LFC across 22 canonical synapse organization transcripts per cluster (WT Vehicle vs 5xFAD Vehicle) showing convergent synaptic gene downregulation (>60% of genes reduced in 5xFAD) spanning cortical excitatory neurons, GABAergic interneurons, astrocytes and the oligodendrocyte lineage. Only clusters with sufficient nuclear representation for single-cell resolution are shown. (C) Violin plots of canonical VIP interneuron transcripts Vip, Reln and Pde7b across WT Vehicle and 5xFAD Vehicle conditions confirming selective transcriptional reduction in the disease state.

Visualization of synapse organization gene expression across clusters revealed convergent downregulation of individual synaptic transcripts across excitatory neurons, GABAergic interneurons and glia (Figure 2B). At the pathway level, GO Biological Process enrichment for synapse organization terms was most significantly concentrated in VIP/Reln interneurons (regulation of synapse structure or activity: 44 genes, padj = 1.5×10^−14; regulation of synapse organization: 43 genes, padj = 1.9×10^−14) while astrocytes, oligodendrocyte lineage clusters and SST interneurons each showed significant enrichment for distinct pathological gene programs including cell-substrate adhesion in astrocytes and glial cell differentiation and calcium ion transport in mature oligodendrocytes. The gene-level convergence in Figure 2B alongside the cluster-level GO specificity together support a model of widespread but cell-type-selective transcriptional disruption in 5xFAD cortex. Reduced expression of canonical VIP interneuron transcripts including Vip, Reln and Pde7b in 5xFAD Vehicle relative to WT Vehicle (Figure 2C) further establishes the VIP interneuron transcriptional signature as a defining feature of the disease landscape in this atlas.

Glial populations exhibited disease-associated transcriptional signatures consistent with established AD pathological states. Astrocytes showed selective loss of cell-substrate adhesion gene programs (padj = 2.4×10^−4, 10 genes) with the receptor-type tyrosine phosphatase Ptprz1 and the pleiotrophic neuroprotective factor Ptn among the most substantially reduced transcripts. Sub-clustering of microglia and astrocytes resolved transcriptionally distinct subpopulations within each glial class (Supplementary Figure 3) and full cluster-level disease signatures across all 16 populations are provided in Supplementary Figure 6.

### VIP/Reln disinhibitory interneurons show pronounced loss of synapse organization gene programs

Among GABAergic interneuron subtypes VIP/Reln interneurons (n = 149 nuclei) showed the most transcriptionally coherent disease-associated disruption. UMAP localization of the three inhibitory subtypes, PVALB (n = 1,820), SST (n = 916) and VIP/Reln (n = 149), is shown in Figure 3A. An excitatory/inhibitory gene heatmap ordered by VIP/Reln LFC (Figure 3B) demonstrates that the most substantial downregulation in 5xFAD VIP interneurons spans transcripts encoding postsynaptic scaffold proteins, ionotropic and metabotropic receptor subunits and intracellular signaling effectors while PVALB and SST interneurons show comparatively diffuse disruption across the same gene panel.

**Figure 3.**
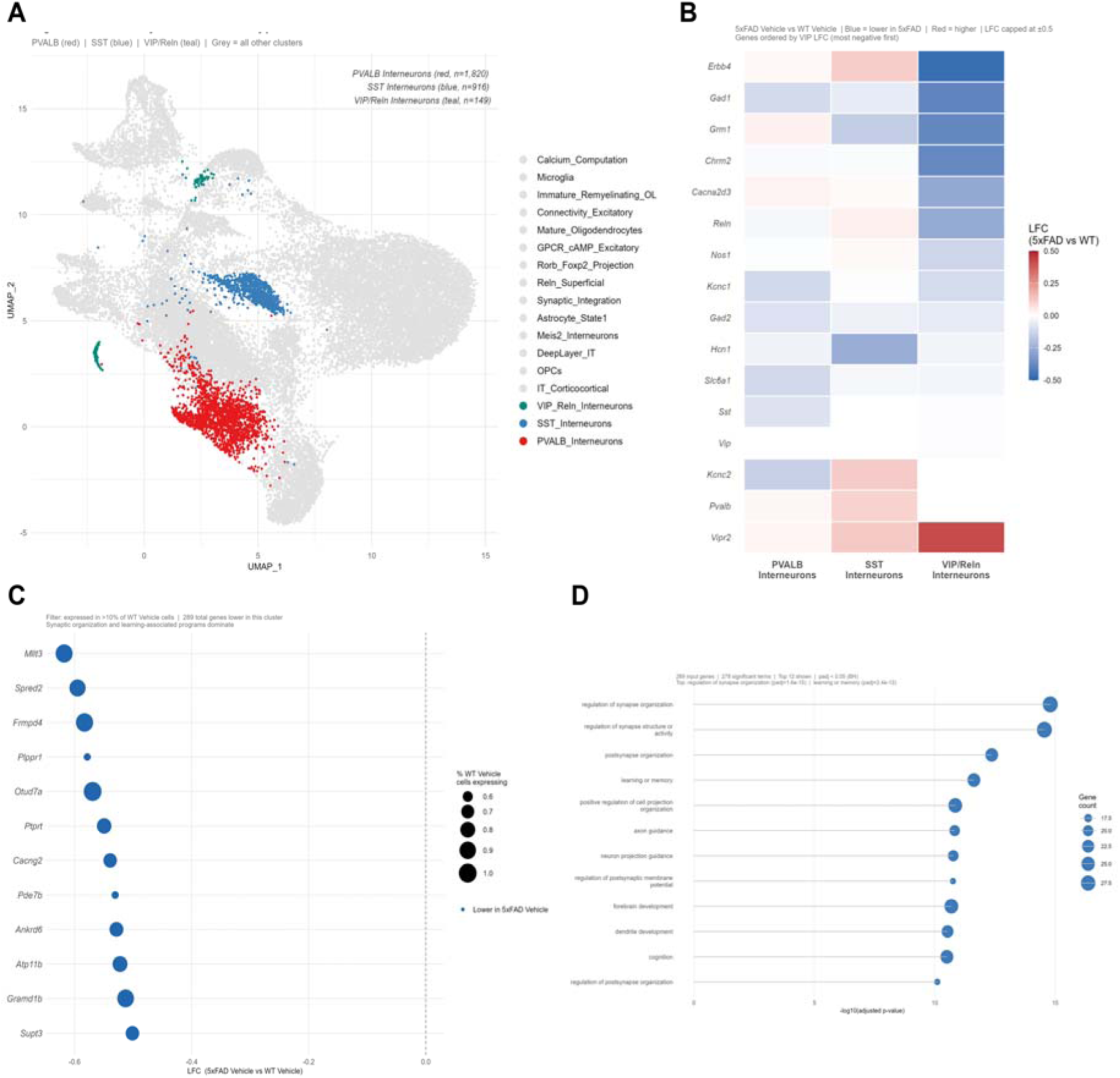
VIP/Reln disinhibitory interneurons show pronounced loss of synapse organization gene programs in 5xFAD. (A) UMAP embedding showing the spatial distribution of PVALB (n = 1,820), SST (n = 916) and VIP/Reln (teal, n = 149) GABAergic interneuron subtypes within the full atlas. (B) Excitatory/inhibitory gene heatmap ordered by VIP/Reln disease LFC where blue encodes transcriptional reduction and red encodes elevation in 5xFAD Vehicle relative to WT Vehicle; LFC capped at +/-0.5. (C) Dot plot of the top transcripts reduced in 5xFAD VIP/Reln interneurons (289 total downregulated genes; WT Vehicle vs 5xFAD Vehicle) where dot size encodes percent nuclei expressing and dot color encodes mean LFC magnitude. (D) GO Biological Process enrichment of disease-downregulated genes in VIP/Reln interneurons showing top terms: regulation of synapse structure or activity (padj = 1.5×10^−14, 44 genes), regulation of synapse organization (padj = 1.9×10^−14, 43 genes), postsynapse organization (padj = 1.1×10^−13, 38 genes) and dendrite development (padj = 6.1×10^−10, 33 genes); Benjamini-Hochberg corrected at padj < 0.05.

Comparing 5xFAD Vehicle (n = 53 nuclei) to WT Vehicle (n = 51 nuclei) 289 candidate genes were reduced in VIP/Reln interneurons with reductions spanning synaptic organization, voltage-gated and ligand-gated calcium signaling and neuropeptide receptor transcripts as confirmed by the top disease-effect gene dotplot (Figure 3C). GO Biological Process enrichment of disease-downregulated genes yielded highly significant pathway-level specificity with top terms including regulation of synapse structure or activity (padj = 1.5×10^−14, 44 genes), regulation of synapse organization (padj = 1.9×10^−14, 43 genes), postsynapse organization (padj = 1.1×10^−13, 38 genes) and dendrite development (padj = 6.1×10^−10, 33 genes) (Figure 3D). The GO enrichment depth of VIP/Reln interneurons substantially exceeded that of PVALB and SST interneurons (Supplementary Figure 7) supporting VIP/Reln interneurons as the primary interneuron finding of this atlas. Glial subcluster UMAPs resolving transcriptionally distinct subpopulations within microglia and astrocytes are presented in Supplementary Figure 3.

### Rolipram treatment is associated with VIP interneuron-enriched transcriptional rescue

Comparing 5xFAD Rolipram to 5xFAD Vehicle across all 16 clusters, VIP/Reln interneurons yielded the strongest true rescue signal. True rescue genes, defined as transcripts reduced in 5xFAD Vehicle relative to WT Vehicle (LFC < −0.25) and restored by rolipram (LFC > +0.25) with expression in >5% of nuclei, were most abundant in VIP/Reln interneurons (n = 72) followed by astrocytes (n = 42) while remaining clusters showed substantially fewer rescued transcripts (Figure 4A), confirming that VIP/Reln interneurons are the preferentially rolipram-responsive inhibitory subtype in this atlas.

**Figure 4.**
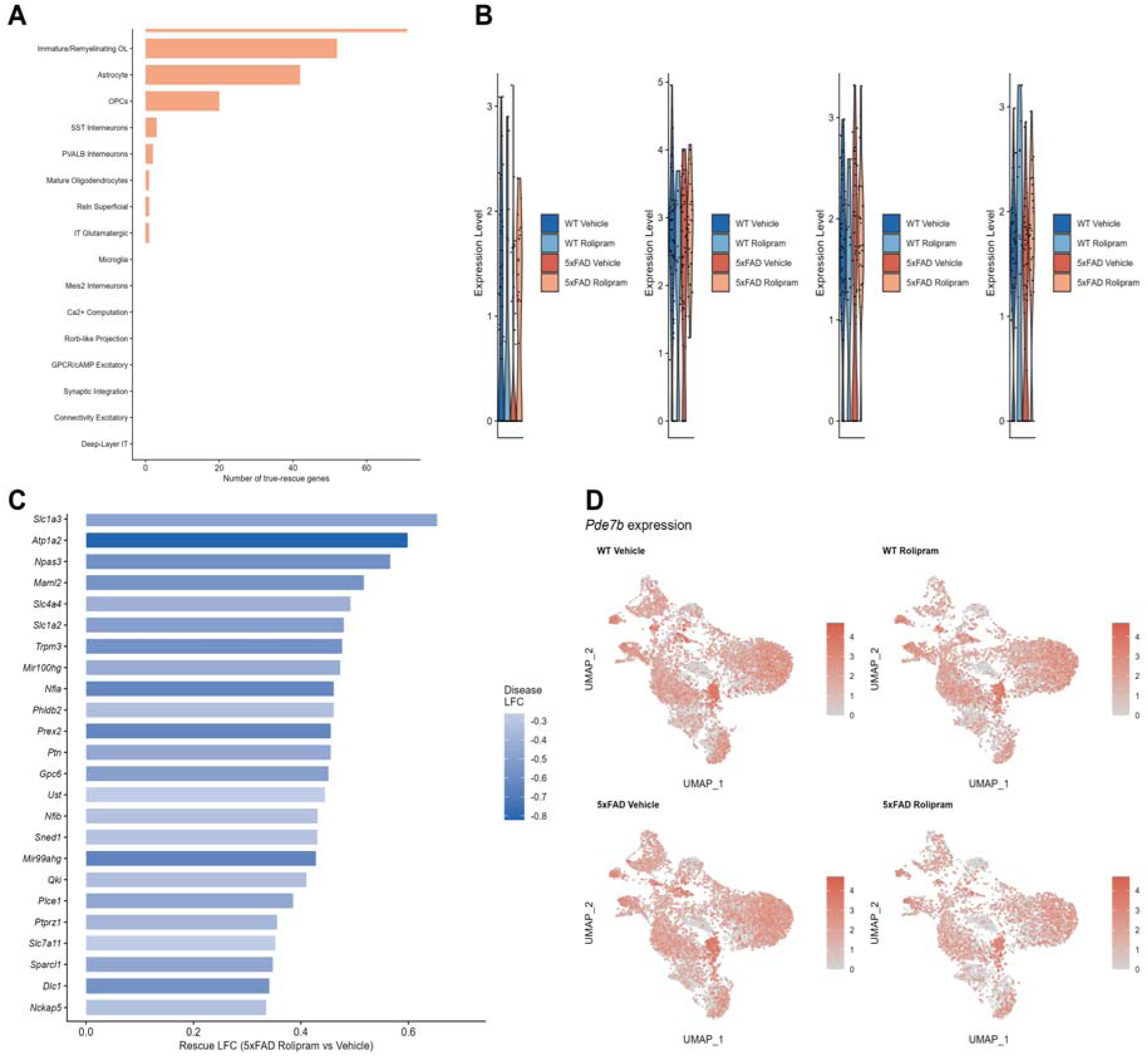
Rolipram treatment is associated with VIP interneuron-enriched transcriptional rescue in 5xFAD. (5xFAD Rolipram vs 5xFAD Vehicle.) (A) Bar chart of true rescue gene counts per cluster (LFC < −0.25 in disease and LFC > +0.25 in rescue, >5% nuclei) showing VIP/Reln interneurons as the most rolipram-responsive population (n = 72) followed by astrocytes (n = 42) with substantially fewer rescued transcripts across remaining clusters. (B) Violin plots of top VIP/Reln true rescue genes including Pde7b across WT Vehicle, 5xFAD Vehicle and 5xFAD Rolipram conditions confirming that rescued expression reflects genuine restoration of disease-suppressed transcription rather than nonspecific pharmacological induction. (C) Astrocyte rescue LFC dot plot of the top 25 rescue genes showing restored glial glutamate transporter expression (Slc1a2, Slc1a3) and suppressed reactive phosphodiesterase transcripts (Pde4d, Pde10a) where dot color encodes disease LFC with blue indicating transcriptional reduction in 5xFAD Vehicle. (D) Pde7b UMAP feature plots across all four experimental groups confirming disease-context-specific transcriptional reduction in 5xFAD Vehicle, restoration in 5xFAD Rolipram and expression in WT Rolipram comparable to WT Vehicle with a consistent color scale across all panels.

Violin plots of top VIP true rescue genes including Pde7b confirm that the rescue effect reflects genuine restoration of disease-suppressed transcription rather than nonspecific pharmacological induction (Figure 4B). In astrocytes rolipram was associated with restored expression of the glial glutamate transporters Slc1a2 and Slc1a3 alongside suppression of reactive phosphodiesterase transcripts Pde4d and Pde10a (Figure 4C) with GO enrichment of astrocyte true rescue genes pointing to partial restoration of homeostatic glial identity through glial cell differentiation (6 genes, padj = 0.004) and gliogenesis (6 genes, padj = 0.013) programs. Pde7b feature plots across all four experimental groups (Figure 4D) confirm selective transcriptional reduction in 5xFAD Vehicle VIP interneurons with restoration in 5xFAD Rolipram and no change between WT Vehicle and WT Rolipram, the defining signature of disease-context-specific pharmacological rescue. The full VIP rescue gene dotplot is provided in Supplementary Figure 8; the absence of significant GO enrichment terms at padj < 0.05 for the 72 VIP true rescue genes indicates that the VIP rescue program represents a distributed restoration of synaptic and signaling transcription rather than the selective reactivation of a discrete functional pathway.

### Pde7b identifies VIP interneuron cAMP signaling as a candidate mechanistic substrate

Among the 72 VIP true rescue genes Pde7b, encoding the cAMP-specific phosphodiesterase 7B, displayed the most compelling bidirectional disease-rescue pattern. Violin plots of Pde7b expression in VIP/Reln interneurons across all four experimental groups (Figure 5A) show baseline expression in WT Vehicle (n = 51 nuclei), transcriptional reduction in 5xFAD Vehicle (n = 53 nuclei, LFC −0.53), restored expression in 5xFAD Rolipram (n = 28 nuclei, LFC +0.54) and expression in WT Rolipram comparable to WT Vehicle, confirming disease-context specificity of the rescue effect. Cross-cluster analysis of Pde7b LFC across all 16 populations (Figure 5B) confirms that the bidirectional disease-rescue pattern is most pronounced and nearly symmetrical in VIP/Reln interneurons while the WT drug effect remains near zero, further supporting disease-context specificity.

**Figure 5.**
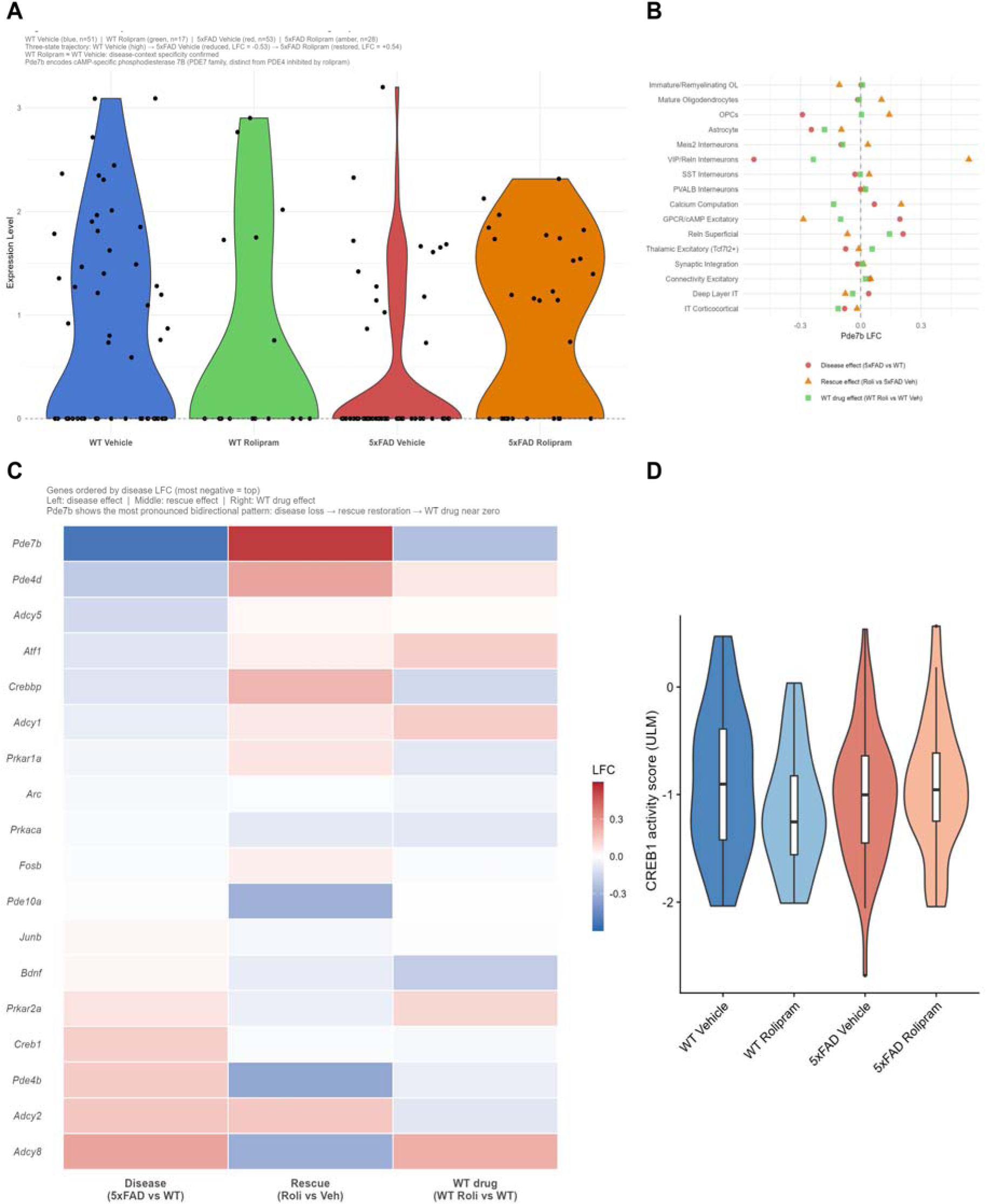
Pde7b and CREB1 identify VIP interneuron cAMP signaling as a candidate mechanistic substrate for rolipram’s procognitive effects. (A) Pde7b expression violin plots in VIP/Reln interneurons across all four experimental groups: WT Vehicle (n = 51), 5xFAD Vehicle (n = 53, LFC −0.53), 5xFAD Rolipram (n = 28, LFC +0.54) and WT Rolipram showing expression comparable to WT Vehicle, confirming disease-context specificity of the pharmacological rescue. (B) Pde7b LFC across all 16 clusters for three pairwise comparisons: disease effect (red), rescue effect (orange) and WT drug effect (green) demonstrating that the most substantial and nearly symmetrical bidirectional pattern is restricted to VIP/Reln interneurons. (C) cAMP pathway gene LFC heatmap in VIP/Reln interneurons showing Pde7b as the most consistently bidirectionally disrupted cAMP pathway component with LFC capped at +/-0.6. (D) decoupleR ULM CREB1 transcriptional activity inference in VIP/Reln interneurons across all four experimental groups showing reduced CREB1 activity in 5xFAD Vehicle and partial restoration in 5xFAD Rolipram supporting a stepwise amyloid-cAMP-PKA-CREB1-Pde7b transcriptional cascade. All results derive from single pooled samples per condition and are hypothesis-generating.

A cAMP pathway gene LFC heatmap in VIP/Reln interneurons (Figure 5C) shows that Pde7b displays the most consistent bidirectional disruption of any profiled cAMP pathway component with other PDE family members and adenylyl cyclase subunits showing smaller or directionally inconsistent effects. These findings support a mechanistic model in which amyloid-driven suppression of intracellular cAMP tone in VIP interneurons reduces PKA-mediated CREB1 phosphorylation at Ser133 which in turn attenuates CREB1-dependent transcription of downstream target genes including Pde7b itself.

To investigate this upstream regulatory mechanism we applied decoupleR (v2.17.0) transcription factor activity inference using the CollecTRI CREB1 target gene network. CREB1 transcriptional activity scores were reduced in 5xFAD Vehicle VIP/Reln interneurons relative to WT Vehicle and partially restored in 5xFAD Rolipram (Figure 5D) supporting the amyloid-driven suppression of cAMP-PKA-CREB1 signaling as the upstream mechanism of Pde7b transcriptional loss and aligning with the established role of CREB1-mediated gene regulation in synaptic consolidation and long-term memory formation [20, 21]. CREB1 transcriptional activity scores across all 16 clusters are provided in Supplementary Figure 4 and glial rescue signatures including a disease-and-rescue LFC heatmap spanning microglia, astrocytes and the oligodendrocyte lineage alongside a cross-cell-type rescue dotplot are presented in Supplementary Figure 9.

Supplementary Figure 2 provides the disease-rescue specificity scatter plot for VIP/Reln interneurons confirming that true rescue genes cluster in the expected bidirectional quadrant when rescue LFC is plotted against disease LFC across the full expressed gene set. Supplementary Figure 5 presents sex-of-origin deconvolution from Xist and Y-linked gene expression confirming largely mutually exclusive male and female nuclear classification within each pooled condition with disease and rescue LFCs for Pde7b, Chrm2 and Reln showing directional consistency across inferred sexes. Supplementary Figure 6 provides GO enrichment for disease-downregulated genes across all 16 clusters confirming that pathway-level synapse organization enrichment is concentrated in VIP/Reln interneurons and contextualizing this finding within the full cellular atlas.

## Discussion

We present a single-nucleus transcriptomic atlas of 5xFAD and wild-type mouse brain under vehicle and rolipram treatment, providing cell-type-resolved characterization of disease-associated transcriptional disruption and its pharmacological modulation across 29,664 nuclei and 16 annotated cell type populations. Profiling excitatory neurons, GABAergic interneuron subtypes, astrocytes, oligodendrocyte lineage cells, OPCs, and microglia within a single 2×2 factorial design enables simultaneous comparison of disease and pharmacological effects across the full cellular composition of the cortex using a consistent LFC framework, a level of inference not accessible from bulk RNA-seq or studies restricted to one or two cell types. Several findings emerge from this atlas that advance the mechanistic understanding of AD cortical pathology and identify cell-type-specific transcriptional substrates for rolipram’s procognitive actions.

Single-nucleus profiling revealed that synaptic gene downregulation in 5xFAD is broadly distributed at the level of individual transcript expression across cell types yet highly concentrated at the GO pathway level within VIP/Reln interneurons. This distinction carries interpretive weight: the shared individual synaptic transcripts reduced across multiple clusters reflect a generalized response, whereas coordinated suppression of synapse organization programs, including regulation of synapse structure or activity (44 genes, padj = 1.5×10^−14) and regulation of synapse organization (43 genes, padj = 1.9×10^−14), is selective to VIP/Reln interneurons and is not reproduced at equivalent statistical depth in PV, SST or excitatory populations. This pattern fits within the broader single-nucleus AD literature: Mathys et al. documented synaptic gene losses as a cardinal transcriptional feature of AD neuronal pathology in both mouse and human cortex (6, 7), and Morabito et al. identified shared chromatin accessibility changes at synaptic loci across cell types in human AD (9). The data suggest that among GABAergic interneuron subtypes, the transcriptional perturbation of synaptic gene programs is selectively concentrated in the disinhibitory VIP interneuron population, a cell type whose pathway-level transcriptional vulnerability has not previously been characterized by GO-level analysis in 5xFAD. Soluble amyloid-beta oligomers, which impair astrocytic glutamate reuptake via excitatory amino acid transporters and disrupt postsynaptic density organization and Hebbian plasticity across neuronal and glial membranes (11, 28), provide a candidate upstream mechanism for the widespread gene-level changes observed in Figure 2B, while the GO-level concentration in VIP/Reln may reflect the particular dependence of this population on activity-dependent regulation of synaptic scaffolds (29).

The identification of VIP/Reln disinhibitory interneurons as the most transcriptionally disrupted inhibitory subtype, with 289 candidate genes lower in 5xFAD and highly significant GO enrichment for synapse organization programs, is consistent with emerging evidence implicating interneuron vulnerability as an early feature of AD pathophysiology (6, 30). Prior transcriptomic studies have characterized PV and SST interneuron vulnerability in AD mouse models and human tissue, with stage-dependent metabolic and synaptic dysregulation in PV interneurons detectable before overt amyloid accumulation and SST interneurons showing region-specific loss in AD (6, 30). These data extend the interneuron vulnerability landscape to the VIP disinhibitory subtype and identify it as the most transcriptionally coherent inhibitory population in 5xFAD cortex at this disease stage. The functional weight of this disruption follows directly from the disinhibitory microcircuit architecture VIP interneurons occupy: selective GABAergic targeting of SST interneurons by VIP interneurons disinhibits SST-mediated dendritic inhibition onto pyramidal neurons thereby gating net excitatory cortical output, such that transcriptional compromise of this population would be expected to reduce cortical gain precisely during the cholinergic and noradrenergic neuromodulatory states that permissively regulate synaptic consolidation and long-term potentiation induction (14, 15). Loss of this disinhibitory gate would attenuate excitatory pyramidal recruitment during the neuromodulatory states that gate Hebbian synaptic plasticity and LTP induction, providing a direct circuit-level mechanistic link between VIP interneuron transcriptional disruption and the spatial memory and contextual fear conditioning deficits observed in 5xFAD mice (16). The relative sparing of PVALB and SST interneurons from coherent GO-level pathway disruption (Supplementary Figure 7) further singles out the VIP population as a candidate early cellular locus of AD inhibitory network pathology.

The glial findings provide independent validation of dataset quality through recovery of established AD pathological signatures with cell-type specificity. Loss of cell-substrate adhesion gene programs in astrocytes (padj = 2.4×10^−4, 10 genes) with selective reduction of the receptor-type tyrosine phosphatase proteoglycan Ptprz1 and the pleiotrophic neuroprotective factor Ptn aligns with the broader literature characterizing AD-associated astrocyte state transitions in which coordinated loss of homeostatic extracellular matrix maintenance and neuroprotective secretory programs occurs alongside transcriptional gain of reactive astrogliosis signatures (35, 36). The absence of canonical A1 neurotoxic astrocyte gene enrichment in 5xFAD astrocytes is consistent with observations in other neurodegeneration contexts (34, 35), supporting the emerging consensus that disease-associated astrocytes adopt context-specific programs that do not map cleanly onto the A1/A2 binary originally defined in neuroinflammation models (35). Both Ptprz1 and Ptn contribute to extracellular matrix organization and neuroprotective signaling, making them specific molecular candidates for the loss of astrocyte homeostatic support in 5xFAD tissue.

The rolipram rescue analysis identifies preferential transcriptional responsiveness of VIP interneurons and astrocytes, with the VIP-enriched signal being both quantitatively dominant and mechanistically coherent. The VIP-preferential rescue is consistent with the enriched expression of VIPR2 (a Gs-coupled VIP receptor generating robust cAMP responses) and high adenylyl cyclase activity in VIP interneurons relative to other subtypes (31). By elevating intracellular cAMP through PDE4 inhibition, rolipram would be predicted to exert its strongest transcriptional effects in cell types with the highest basal cAMP turnover and most active CREB-dependent gene expression, conditions that are characteristic of VIP interneurons given their enriched VIPR2-Gs-adenylyl cyclase signaling axis. The astrocyte rescue signal (n = 42 true rescue genes with GO enrichment for glial cell differentiation and gliogenesis at padj = 0.004 and 0.013, respectively) is consistent with partial restoration of homeostatic astrocyte transcriptional identity, in keeping with evidence that cAMP-PKA signaling promotes the reversion of reactive astrocytes toward a homeostatic gene expression state through CREB-dependent suppression of NF-kB-driven neuroinflammatory transcription (32). Restoration of the glial glutamate transporters Slc1a2 and Slc1a3 alongside glial differentiation transcripts within the astrocyte rescue program suggests rolipram may simultaneously suppress reactive astrogliosis and restore the glutamatergic buffering and neuroprotective secretory functions that pathological reactivity compromises. The absence of significant GO enrichment within the 72 VIP true rescue genes, by contrast, indicates that the VIP rescue program reflects a distributed, pleiotropic restoration of synaptic and signaling transcription rather than the focused reactivation of a discrete functional pathway, a pattern consistent with CREB1 functioning as a master activity-dependent transcriptional regulator whose PKA-mediated reactivation simultaneously de-represses a broad repertoire of downstream synaptic target genes.

Pde7b represents the most direct mechanistic lead generated by this dataset. PDE7B is a high-affinity cAMP-specific phosphodiesterase with selective neural enrichment and regional expression concentrated in cortex, hippocampus, and striatum (33). Its disease-context-specific transcriptional reduction in 5xFAD VIP interneurons followed by rolipram-mediated restoration with WT Rolipram expression comparable to WT Vehicle (Figure 5A) supports a CREB1-dependent transcriptional negative feedback model in which sustained cAMP-PKA signaling under homeostatic conditions drives CREB1-mediated Pde7b transcription as an autoinhibitory brake on cAMP accumulation. In 5xFAD VIP interneurons, amyloid-driven reduction of intracellular cAMP tone attenuates PKA activity and CREB1 phosphorylation at Ser133, thereby suppressing Pde7b transcription, while rolipram restores cAMP to a level sufficient to reactivate CREB1-dependent Pde7b expression exclusively in the disease context where cAMP is pathologically depleted rather than in WT interneurons operating at homeostatic cAMP levels. Disease-context specificity carries mechanistic significance: it predicts that PDE4 inhibition normalizes rather than supraphysiologically amplifies cAMP-PKA signaling in VIP interneurons, thereby reducing the likelihood of excitotoxic or off-target overstimulation. decoupleR-based CREB1 activity inference, showing attenuated transcriptional activity in 5xFAD vehicle VIP interneurons with partial restoration following rolipram treatment (Figure 5D), supports this model and positions Pde7b within a coherent upstream regulatory cascade: amyloid pathology reduces cAMP tone, attenuates PKA-mediated CREB1 phosphorylation, suppresses CREB1-dependent Pde7b transcription, and rolipram reverses this cascade through PDE4 inhibition upstream. Direct validation will require quantification of PDE7B protein by immunofluorescence in VIP interneuron-identified cells, intracellular cAMP measurement in FACS-isolated VIP interneurons from treated and untreated 5xFAD mice and functional correlation of Pde7b restoration with CREB1 phosphorylation and downstream target gene re-expression, all of which are currently in progress.

Several important limitations must be acknowledged. Most critically, each experimental condition is represented by a single pooled biological sample containing one male and one female animal. Without biological replication at the sample level, standard inferential statistical frameworks are not applicable, and all LFC values are descriptive comparisons rather than statistically tested estimates (25). This is an inherent constraint of the pilot study design, chosen to enable atlas-scale profiling across all four conditions simultaneously before scaling to a fully powered cohort. The 5xFAD Rolipram condition yielded fewer nuclei (n = 4,913) than the other three conditions (n = 7,100-8,840), which reduces statistical power for rescue analyses in small clusters and should be considered when comparing rescue magnitudes across cell types. The whole-hemisphere dissociation approach yields a predominantly cortical atlas with a minor thalamic contribution, leaving hippocampal and striatal cell populations unrepresented despite their central roles in AD-associated episodic memory and corticostriatal circuit dysfunction, respectively, and both regions should be prioritized in subsequent profiling efforts. Sex-of-origin deconvolution using Xist and Y-linked gene expression (Supplementary Figure 5) provides partial mitigation of the pooling limitation, with disease and rescue LFCs for Pde7b, Chrm2 and Reln showing directional consistency across inferred male and female nuclear fractions, supporting the internal reproducibility of these transcriptional effects within the constraints of the pilot design. All findings remain hypothesis-generating and require validation in independently powered, biologically replicated cohorts with sufficient statistical resolution for formal differential expression testing.

## Conclusion

Single-nucleus transcriptomics of 5xFAD mouse brain under rolipram treatment reveals a layered picture of AD cortical pathology and its pharmacological modulation. At the disease level transcriptional disruption is cell-type-specific: while individual synaptic transcripts are broadly reduced across cortical cell populations, coherent GO-level pathway disruption of synapse organization programs is selectively concentrated in VIP/Reln disinhibitory interneurons, identifying this population as the most transcriptionally vulnerable GABAergic subtype in 5xFAD cortex at this disease stage. Glial populations independently validate the atlas through recovery of an established AD astrocyte signature characterized by loss of cell-substrate adhesion and neuroprotective gene programs with cell-type specificity consistent with the broader AD neurogenomics literature.

At the pharmacological level rolipram’s transcriptional rescue is preferential rather than global, with VIP interneurons and astrocytes accounting for the dominant rescue signal and the VIP rescue mechanistically grounded in the Pde7b/CREB1 regulatory axis. The disease-context specificity of Pde7b rescue, restored in 5xFAD Rolipram and unaltered in WT Rolipram, supports a cAMP transcriptional negative feedback model in which PDE4 inhibition normalizes pathologically suppressed cAMP-PKA-CREB1 signaling selectively in cell types where amyloid pathology has disrupted cAMP homeostasis. decoupleR CREB1 activity inference supports this model and identifies CREB1 as an accessible transcription factor node for downstream mechanistic dissection and targeted therapeutic intervention.

As a pilot study all findings are descriptive and hypothesis-generating. The experimental priorities arising from this dataset are: validation of PDE7B protein reduction in VIP interneurons by immunofluorescence in independently powered and biologically replicated cohorts; functional demonstration that Pde7b transcriptional restoration correlates with recovered CREB1 phosphorylation at Ser133 and re-expression of downstream synaptic target genes; behavioral characterization of VIP interneuron-specific conditional Pde7b rescue or viral overexpression in 5xFAD mice; and extension of the atlas to hippocampal and striatal cell populations in a fully replicated experimental design. Together, this dataset suggests a cell-type-resolved transcriptional foundation for mechanistic studies of rolipram’s procognitive action and identifies VIP interneuron cAMP-PKA-CREB1 signaling as an accessible and disease-context-specific cellular substrate for therapeutic intervention in AD.

## Materials and Methods

### Animals

5xFAD transgenic mice (B6SJL-Tg(APPSwFlLon,PSEN1*M146L*L286V)6799Vas/Mmjax) on a B6/SJL mixed background and wild-type littermate controls were used. Each pooled experimental sample contained brain tissue from one male and one female animal aged 6.5-8 months at Alzet osmotic pump implantation. Brains were harvested after 30 days of continuous treatment (animal age 7.5-9 months at harvest). All animal procedures were approved by the IACUC of Texas Tech University Health Sciences Center (protocol # 23014) and performed in accordance with the approved protocol and NIH guidelines.

### Rolipram drug formulation and Alzet osmotic pump implantation

Rolipram (MedChemExpress, CAT# HY-16900, Lot 278297) was prepared at 10 µg/µL in a vehicle of 45% DMSO, 40% PEG300, 13% saline, and 2% Tween 80. Dose was calculated individually for each animal from its measured body weight to achieve a target of 0.03 mg/kg/day. Alzet micro-osmotic pumps (Model 1004, DURECT Corporation; nominal delivery rate 0.11 µL/hour, 100 uL reservoir) were loaded with the rolipram or vehicle solution and primed by incubation in sterile 0.9% saline at 37°C for 48 hours prior to implantation, per manufacturer instructions. Mice were anesthetized with isoflurane for subcutaneous pump implantation between the scapulae.

### Tissue collection and nuclei isolation

After 30 days of treatment, mice were transcardially perfused with ice-cold 1x PBS containing heparin. Brains were hemisected on ice. The right hemisphere was used for single-nucleus RNA sequencing and the left hemisphere was post-fixed in 4% paraformaldehyde. The right hemisphere was snap-frozen and stored at −80°C until nuclei isolation. Nuclei were isolated from frozen right-hemisphere tissue using the S.O.F. OptiPrep discontinuous density gradient protocol. Briefly, frozen tissue was thawed and dissociated in ice-cold Nuclei Extraction Buffer (Miltenyi Biotec) using a gentleMACS Octo Dissociator, followed by density gradient centrifugation on a 25-30-35% iodixanol step gradient at 2,486 x g for 41 min at 4 degrees C. The white nuclei band at the 30-35% interface was collected and washed twice. Nuclei concentration and integrity were assessed by LUNA-FL using the AO/PI Cell Viability Kit. Immediately prior to fixation, nuclei from one male and one female animal were pooled per condition (n = 1 pooled biological sample per condition, n = 2 animals per condition, n = 8 animals total).

### Single-nucleus RNA sequencing library preparation

Single-nucleus RNA sequencing libraries were prepared using the Parse Biosciences Evercode WT v3 kit according to the manufacturer’s protocol. The Evercode WT v3 platform employs split-pool combinatorial barcoding in fixed and permeabilized nuclei. Four experimental samples were processed: WT Vehicle, WT Rolipram, 5xFAD Vehicle, and 5xFAD Rolipram. Libraries were submitted to the Next Generation Sequencing Core Facility at Oklahoma State University for quality control and Illumina library preparation, then sent to Parse Biosciences for sequencing.

### Single-nucleus data processing and quality control

Raw FASTQ files were processed using the Parse Biosciences Trailmaker pipeline, which aligned reads to the mouse reference genome, and generated nucleus-by-gene count matrices. The dataset was further processed using Seurat v5.5.0 in R v4.6.0. Following quality control, 29,664 nuclei were retained across four conditions. Unsupervised graph-based clustering identified 24 transcriptional clusters, consolidated to 16 biologically annotated populations by merging clusters sharing identical marker gene profiles. Cell type identities were assigned by manual inspection of top marker genes cross-referenced against published mouse cortex atlases (26, 27).

### Differential expression and log fold change analysis

Log fold changes (LFC) were computed as the difference in mean log-normalized expression between condition groups within each cluster. Three comparisons were performed: disease effect (5xFAD Vehicle vs WT Vehicle), rescue effect (5xFAD Rolipram vs 5xFAD Vehicle) and WT drug effect (WT Rolipram vs WT Vehicle). Candidate differentially expressed genes were defined by |LFC| > 0.25 and expression in more than 5% of nuclei. Artifact genes were excluded prior to analysis, encompassing mitochondrial transcripts (mt-prefix), ribosomal subunits (Rpl/Rps), pseudogenes (Gm+number, digit+Rik) and other high-abundance non-informative transcripts (Malat1, Xist, Cdk8, Rsrp1, Cfap54, Rbm25). True rescue genes were defined by LFC < −0.25 in the disease comparison and LFC > +0.25 in the rescue comparison with expression in more than 5% of nuclei. All LFC values represent descriptive statistics; the absence of biological replication precludes standard differential expression testing and adjusted p-values are not reported for between-condition comparisons (25).

### Sex-of-origin inference

Per-nucleus sex was inferred from expression of Xist (female-specific) and the Y-linked genes Ddx3y, Eif2s3y, and Uty (male-specific). Nuclei with detectable Xist and no detectable Y-linked gene expression were classified as female; nuclei with no detectable Xist and detectable Y-linked gene expression were classified as male; all other nuclei were classified as ambiguous and excluded from sex-stratified analyses. Disease and rescue effect LFCs for Pde7b, Chrm2, Reln, and Vipr2 were computed separately within inferred-male and inferred-female VIP/Reln interneurons to assess directional consistency (Supplementary Figure 5).

### Gene Ontology enrichment analysis

GO Biological Process enrichment was performed using clusterProfiler v4.20.0 [Wu et al., 2021] with org.Mm.eg.db v3.19.1. Benjamini-Hochberg correction was applied with padj < 0.05. Enrichment was performed on disease-downregulated genes (LFC < −0.25, >5% nuclei) per cluster and on the VIP/Reln true rescue gene set. Full GO enrichment results for all clusters are provided in Supplementary Table 5.

### Glial sub-clustering

Microglia and astrocyte sub-clustering was performed by subsetting each population and re-running feature selection (top 2,000 variable genes), PCA, and graph-based clustering at resolution 0.2. UMAP was computed on the sub-clustered population for visualization. Sub-cluster UMAPs are provided in Supplementary Figure 3.

### Transcription factor activity inference

Transcription factor activity inference was performed using decoupleR v2.17.0 [Badia-i-Mompel et al., 2022] with the CollecTRI CREB1 target gene regulatory network. The Univariate Linear Model (ULM) method was applied to the log-normalized expression matrix of VIP/Reln interneurons (n = 149 nuclei). Mean CREB1 activity scores were compared between WT Vehicle (n = 51) and 5xFAD Vehicle (n = 53) nuclei. CREB1 activity across all 16 clusters is provided in Supplementary Figure 4; per-cell activity scores for all 680 assessed transcription factors across all clusters and conditions are provided in Supplementary Table 6.

## Author Contributions

A.P.: conceptualization, animal husbandry, osmotic pump surgery, tissue collection, nuclei isolation, single-nucleus library preparation, bioinformatic analysis, data visualization, writing (original draft). A.D.: animal husbandry, osmotic pump surgery, tissue collection, nuclei isolation, single-nucleus library preparation. H.W.: conceptualization, funding acquisition, supervision, writing (review and editing). All authors reviewed and approved the final manuscript.

## Acknowledgements

The authors thank the Next Generation Sequencing Core Facility at Oklahoma State University for library preparation services, and Parse Biosciences for sequencing and Trailmaker platform support. This work was supported by NIH/NIA grant AG072510. The authors declare no competing interests.

