## Supplementary figures and images for "Single-nucleus RNA-seq atlas of 5xFAD mouse brain maps cell-type-specific transcriptional disruption and preferential rolipram rescue in VIP interneurons"

### SuppFig1_composite.png

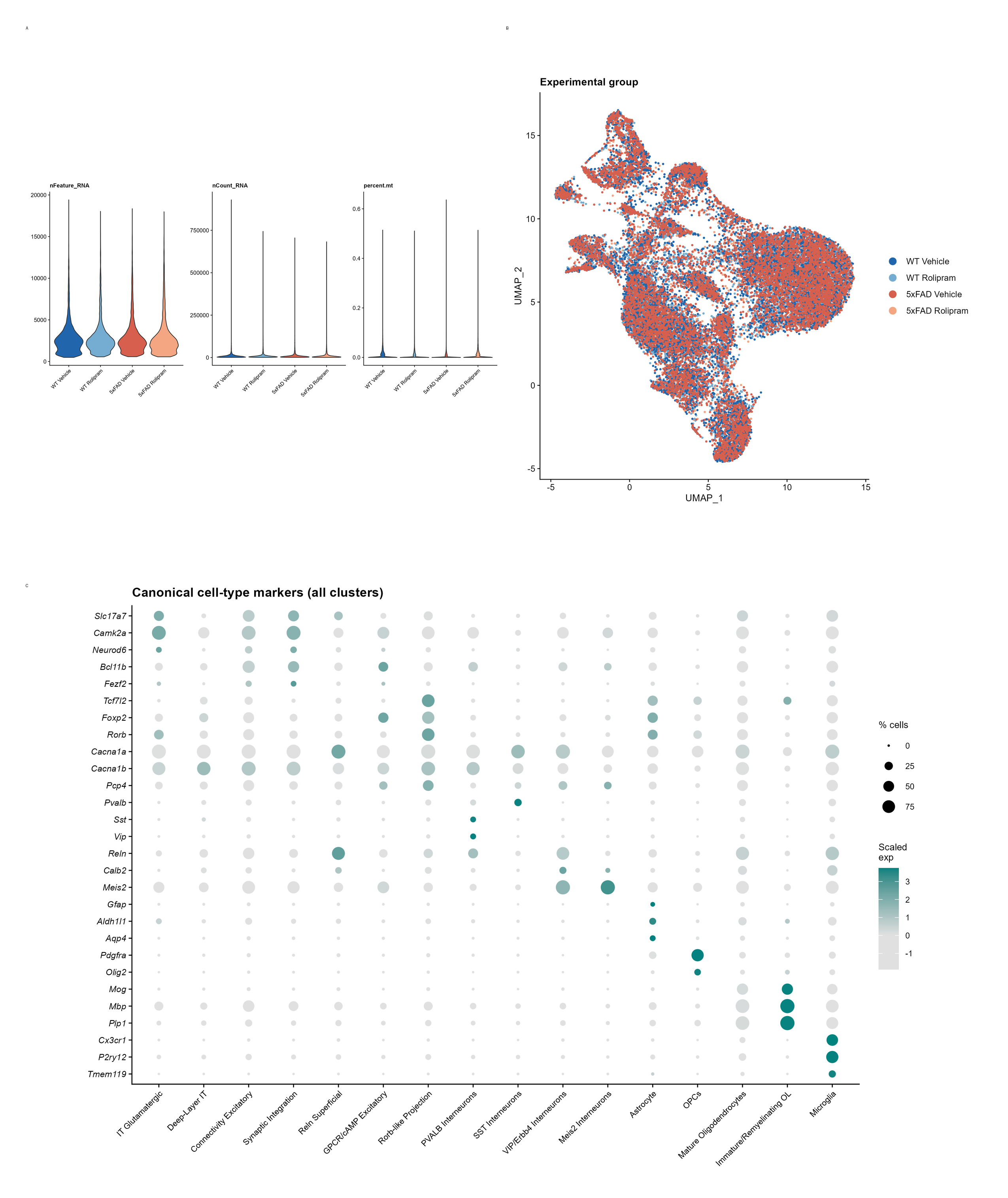

### SuppFig2_composite.png

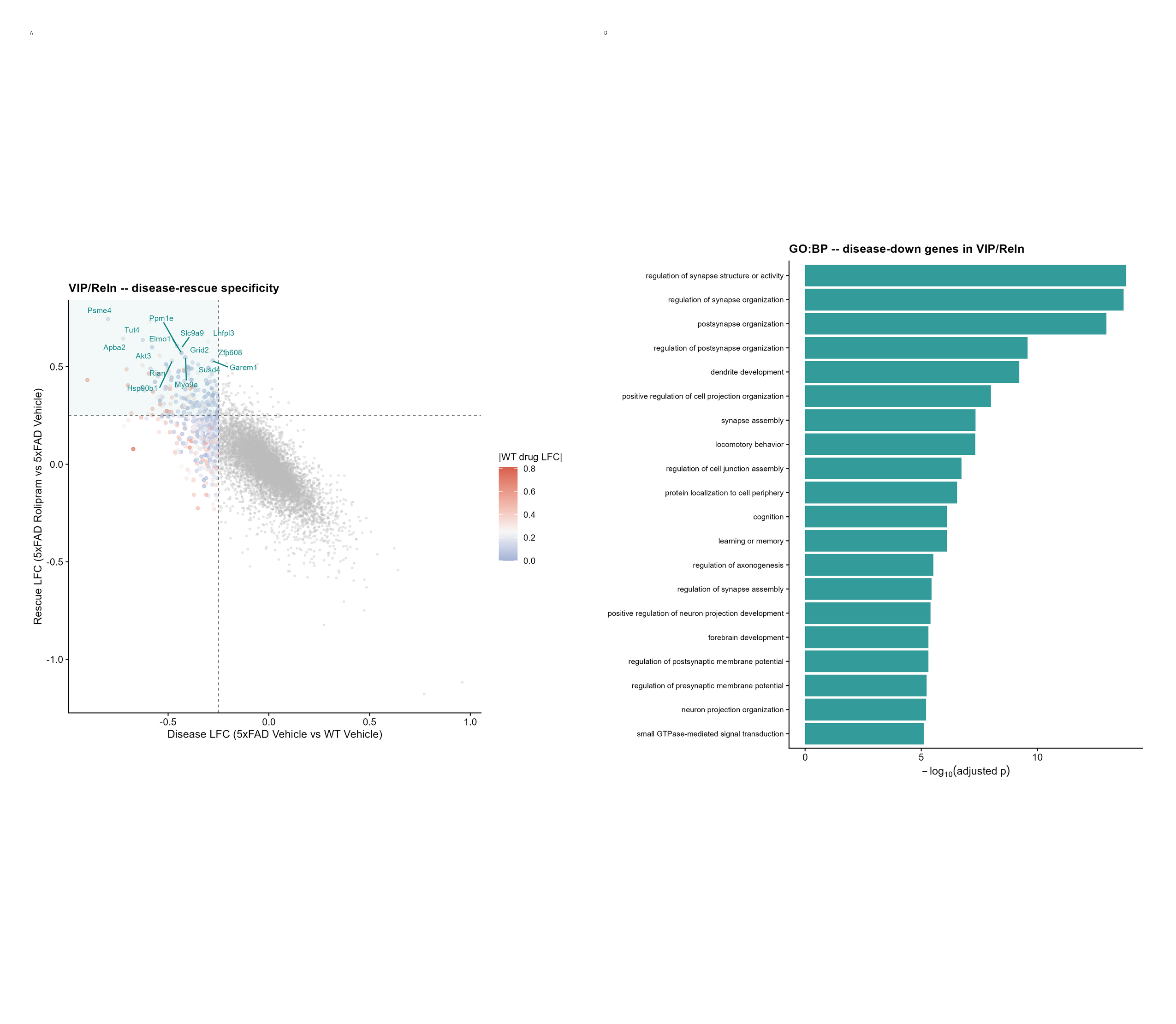

### SuppFig3_composite.png

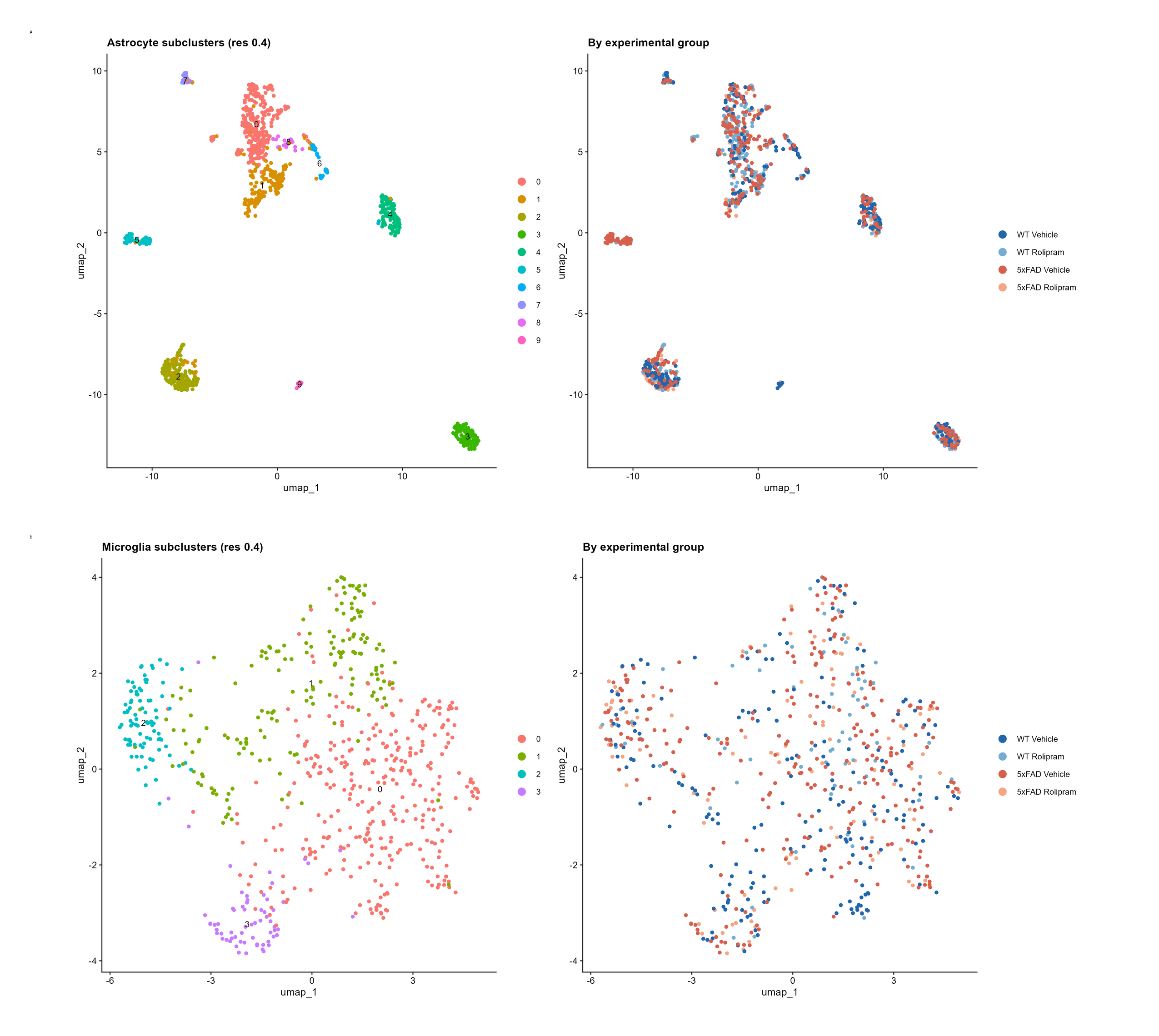

### SuppFig4_composite.png

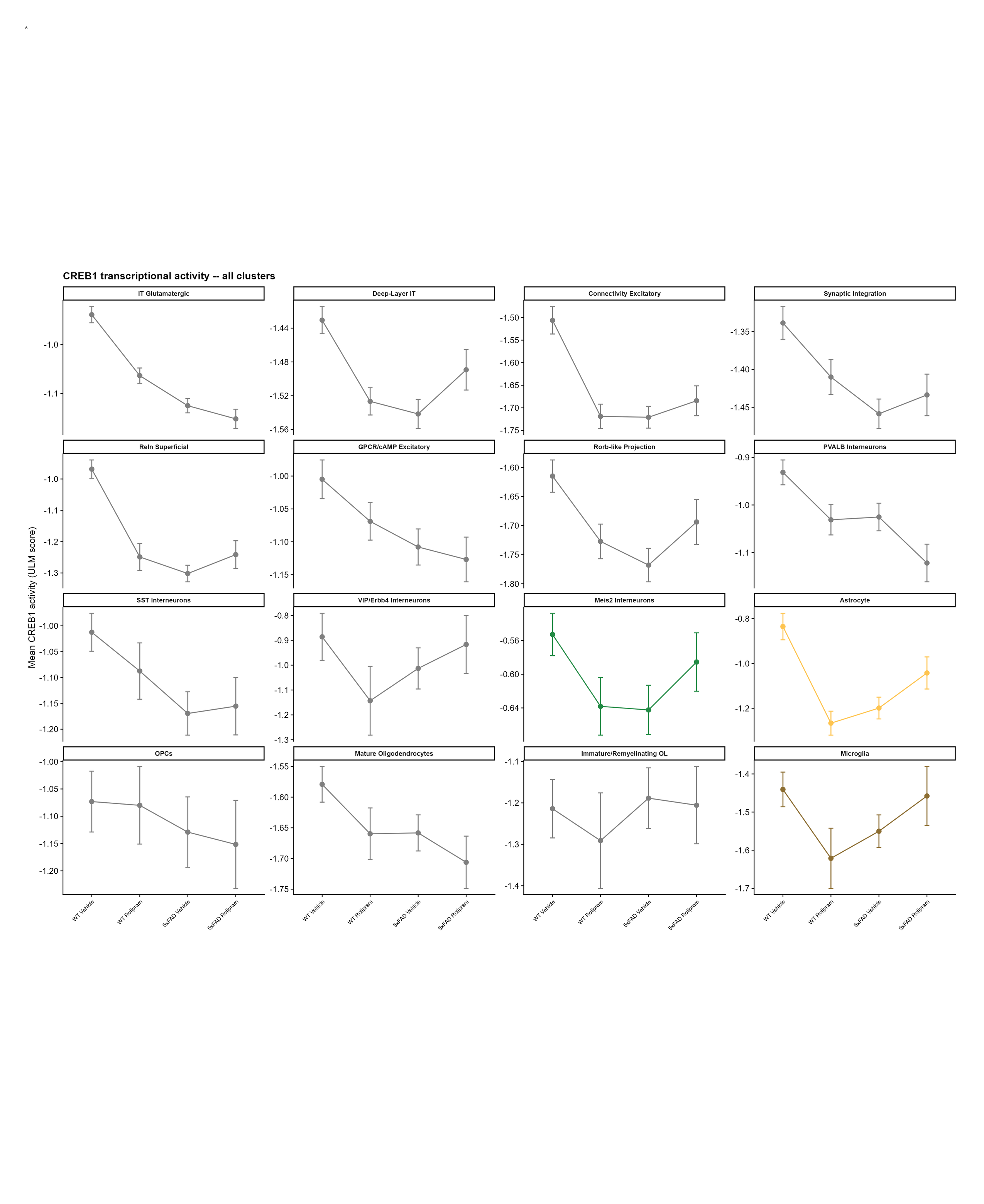

### SuppFig5_composite.png

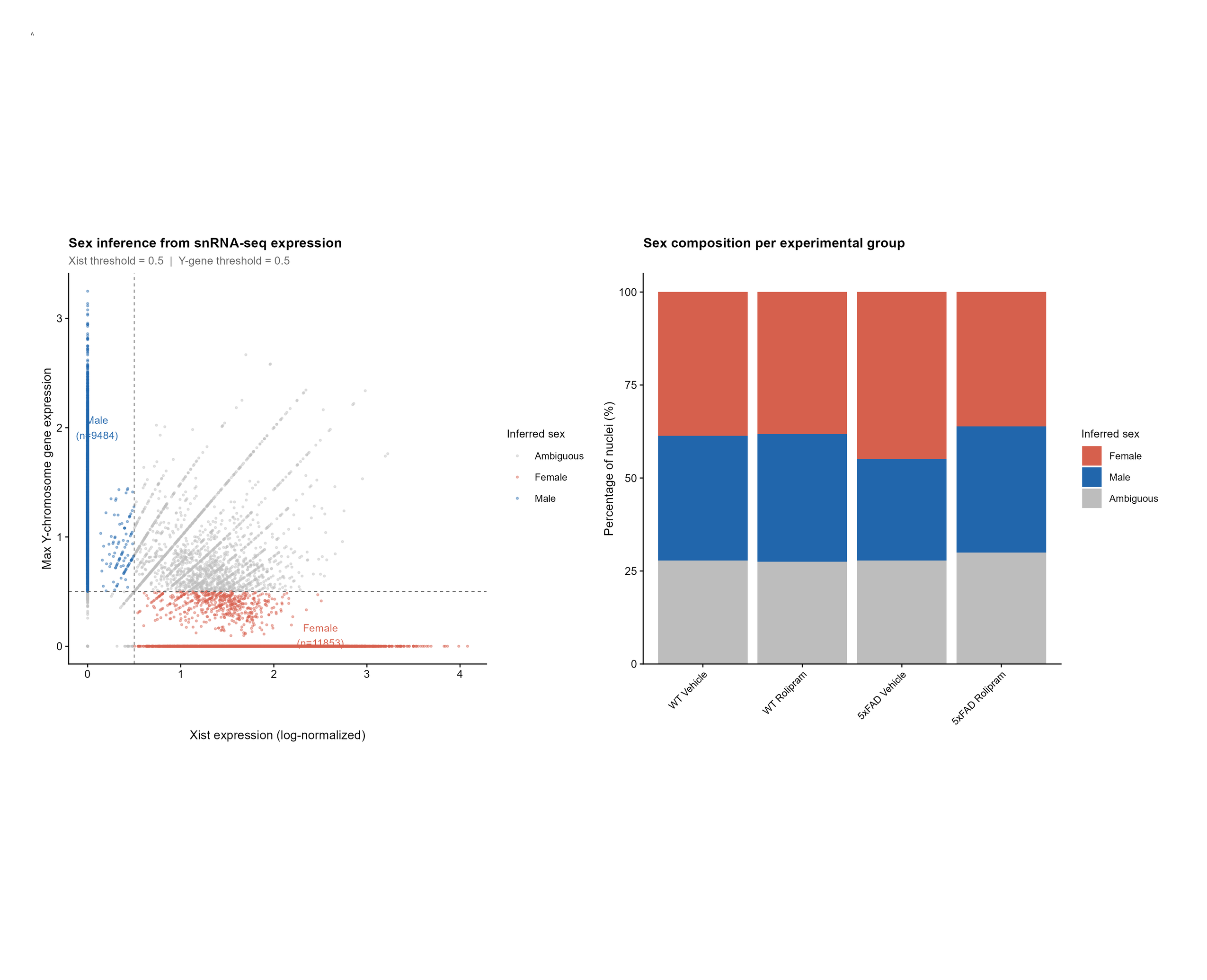

### SuppFig6_composite.png

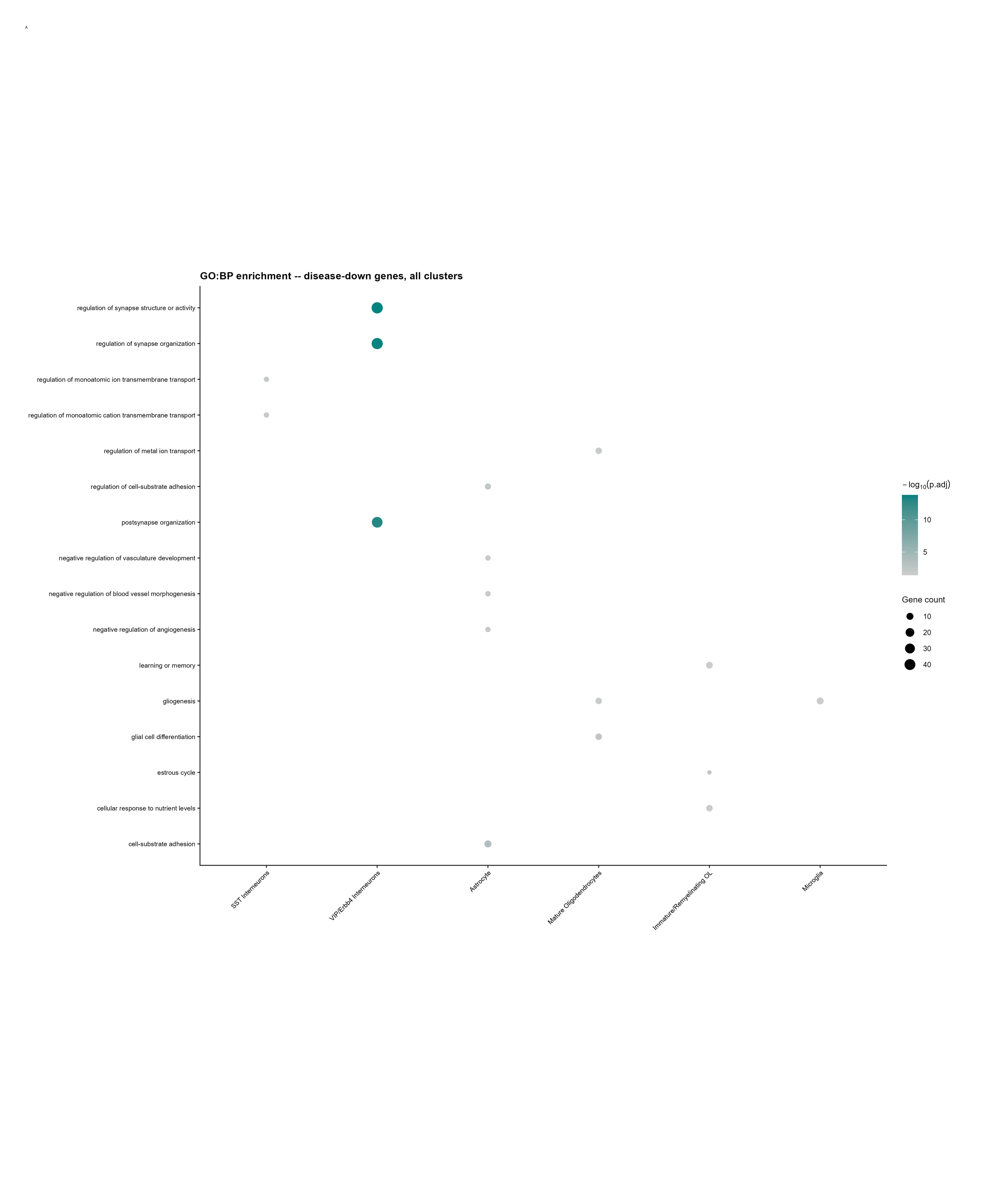

### SuppFig7_composite.png

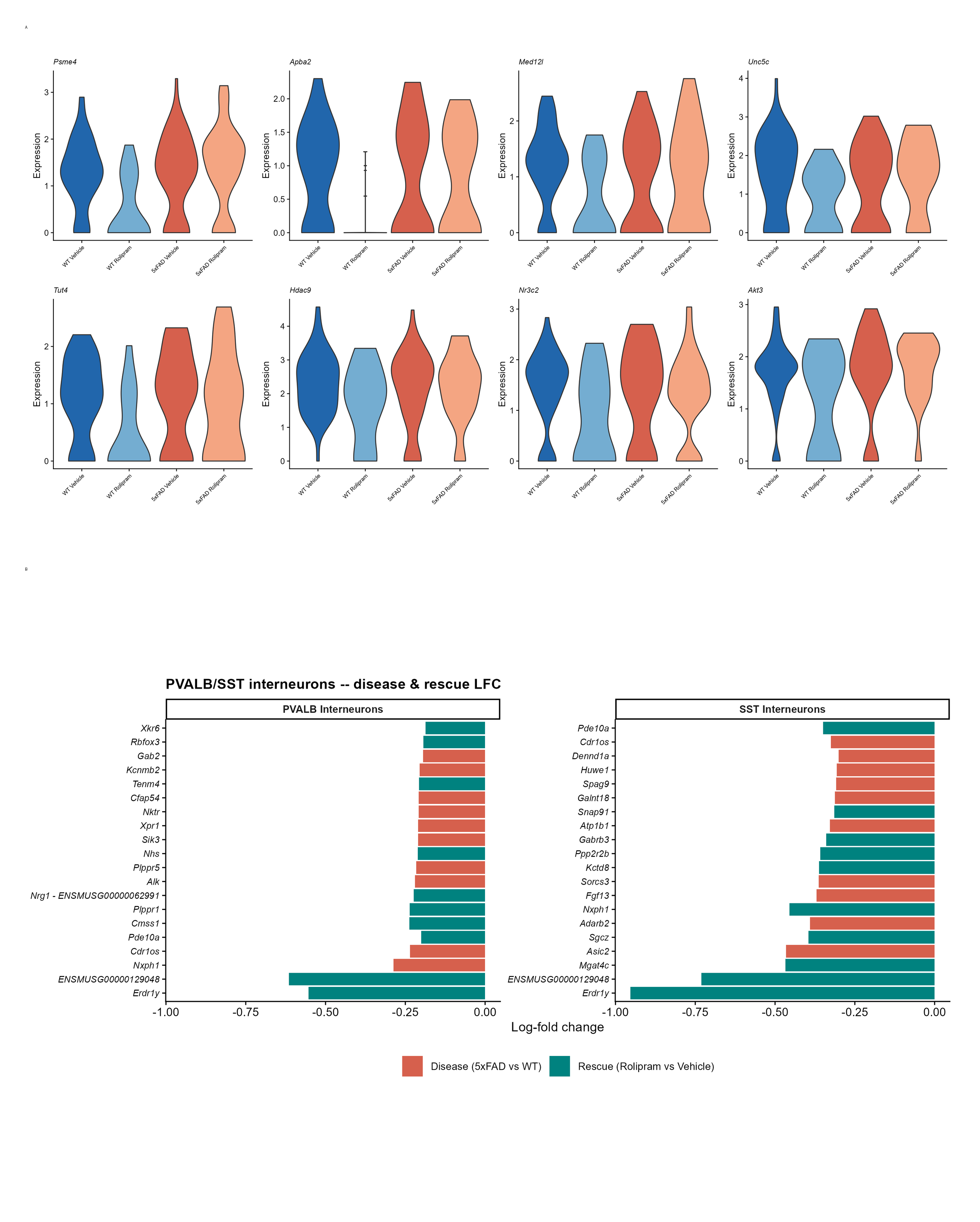

### SuppFig8_composite.png

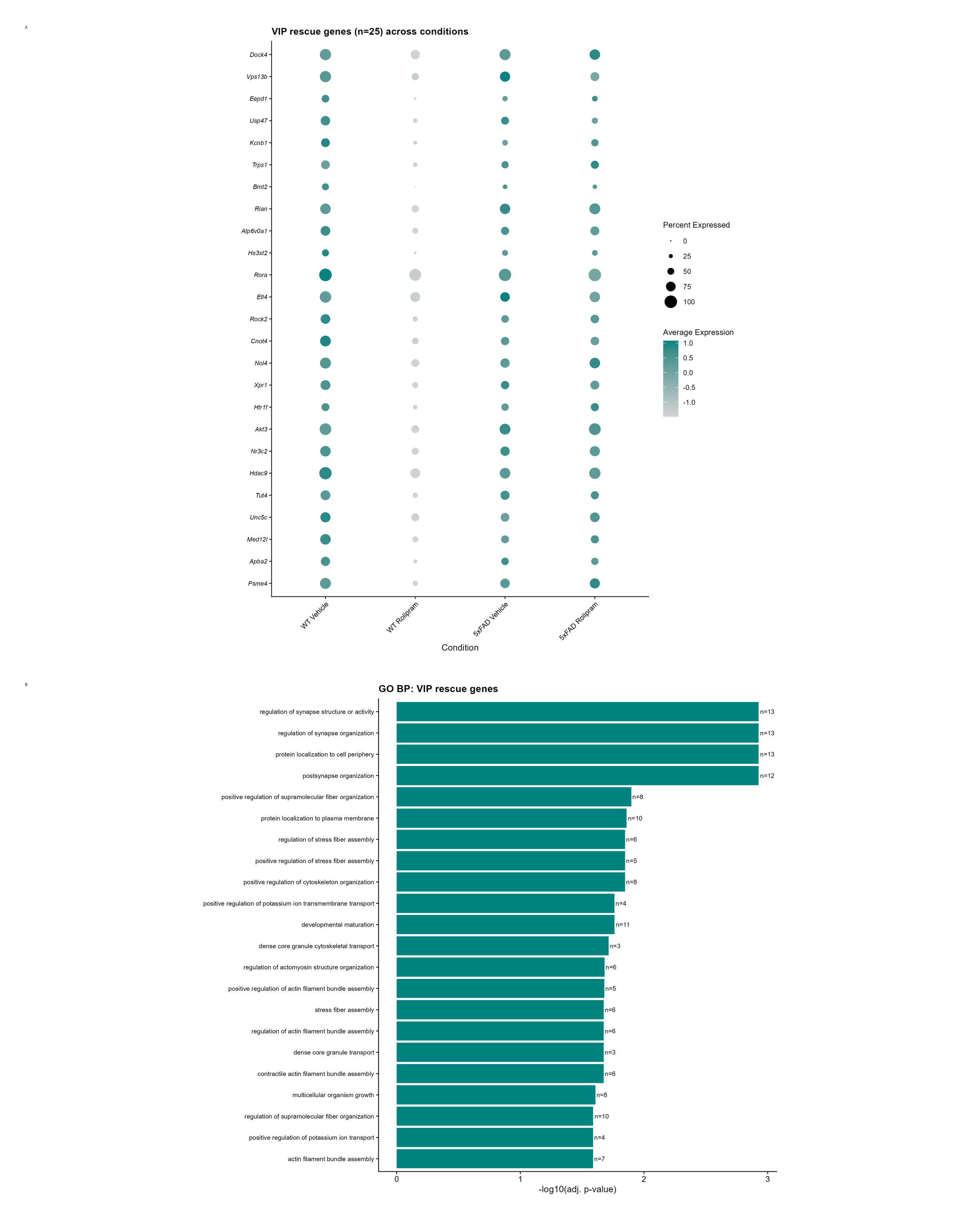

### SuppFig9_composite.png

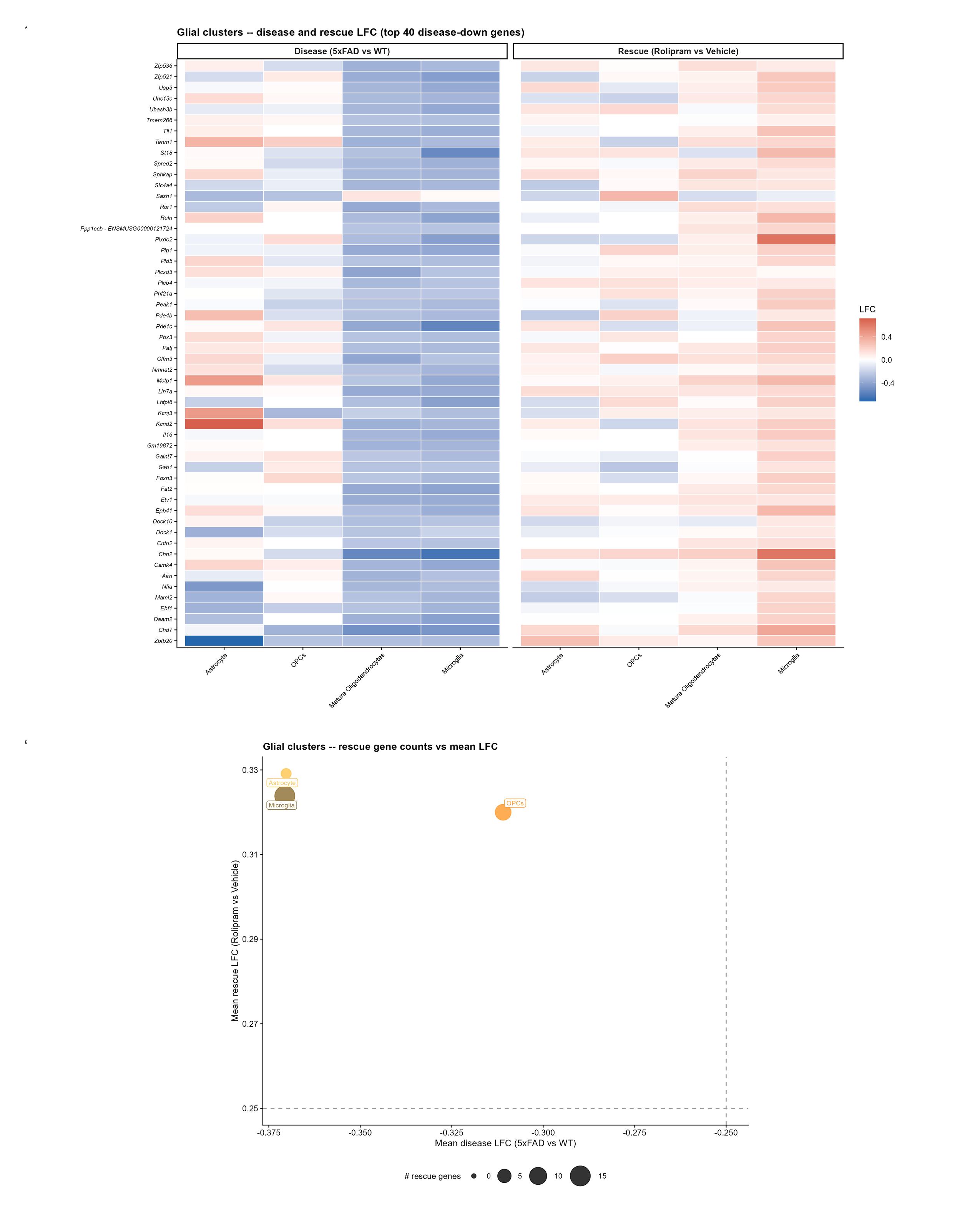
